# Cell-type Specific Alteration of *Dicer1* Accelerates Tumor Progression in Mouse Models of KRAS-driven Lung Adenocarcinoma

**DOI:** 10.64898/2026.05.29.728740

**Authors:** Julie Wells, Richard S. Maser, Rosalinda Doty, Ashley Tucker, Wendy Memishian, Teresa McGee, Naomi Mitchell-Hutchinson, Paige J.K. Ramkissoon, Simon Lesbirel, Jeremy R. Charette, Heidi Munger, Travis Beckett, Carol J. Bult

## Abstract

MicroRNAs (miRNAs) have been widely implicated in cancer initiation and progression, yet examination of the effects of global miRNA disruption on these processes has been limited. We developed novel genetically engineered mouse models of *Kras*-driven pulmonary adenocarcinoma (LUAD) with cell-type-specific disruption of miRNA biosynthesis via *Dicer1* allele deletion, which exhibit significant differences in tumor progression rates and expected survival. *Dicer1* is an RNase III enzyme that is required for the biogenesis of mature, functional miRNAs. Lung tumor progression was accelerated, and expected survival was decreased only when we initiated tumors and deleted one allele of *Dicer1* in club cells and mutated *Dicer1* in alveolar type 2 (AT2) cells. Reversing the cell types by inducing tumorigenesis, deleting one *Dicer1* allele in AT2 cells, and mutating *Dicer1* in club cells modestly accelerated tumor progression and had no effect on expected survival. Collectively, our results demonstrate that *Dicer1* disruption accelerates lung cancer progression in a cell-type-dependent and non-cell-autonomous manner, and our mice represent tools for investigating the roles of miRNAs and miRNA-mediated intercellular communication in tumor progression.

**Summary:** *Kras*-driven mouse models show that *Dicer1* mutations accelerate lung adenocarcinoma (LUAD) progression in a cell-type-dependent manner and suggest that the influence of miRNA-mediated intercellular communication is unidirectional and non-cell-autonomous.

## INTRODUCTION

Lung cancer is the leading cause of cancer deaths worldwide (Siegel et al., 2024, Ferone et al., 2020). Despite recent advances in computed tomography (CT) screening, targeted therapies, and immunotherapies, coupled with increased access to those therapies, the five-year relative survival rate for lung cancer patients is only 28% (Kratzer et al., 2024)(reviewed in (Martinez-Espinosa et al., 2023). The high mortality rate of lung cancer is primarily attributed to metastasis, and most patients present with either regional or distant disease at the time of diagnosis (Drusbosky and Cogle, 2020), (Kratzer et al., 2024). Even patients presenting with localized (stage I) lung cancer who undergo surgery often develop distant metastasis within five years (Rajaram et al., 2024) (reviewed in (Alduais et al., 2023). Collectively, these data suggest that dissemination of cancer cells occurs early during disease progression and that a detailed understanding of the earliest stages of metastasis is needed for effective treatment and management of lung cancer.

Mouse models provide an invaluable opportunity to study the earliest steps in lung cancer progression as the timing and location of tumor induction can be precisely controlled and the entire process, including interactions between tumor and host cells, can be interrogated. Unfortunately, many genetically engineered mouse models (GEMMs) of lung cancer take a long time to develop metastases, have low penetrance and do not metastasize to all of the same sites as in humans (reviewed in (Hebert et al., 2023, Giacobbe and Abate-Shen, 2021). For example, 40% of pulmonary cancers are classified as lung adenocarcinomas (LUADs) and these tumors frequently harbor oncogenic activating mutations in *KRAS* (HGNC:6407) and alterations in the tumor suppressor gene *TP53* (HGNC:11998) *(Zhang et al., 2023, Gillette et al., 2020)*. A widely studied mouse strain for LUAD research is the *Kras^tm4Tyj/+^*;*Trp53^tm1Brn/tm1Brn^*(MGI:3716968) strain, commonly referred to as KP. Following Cre recombinase exposure, KP mice conditionally express oncogenic *Kras*^G12D^ and delete the floxed exons 2-10 from both *Trp53* (Jackson et al., 2005) alleles. While pulmonary alveolar type II (AT2; CL:0002063) cells are widely regarded as the cell of origin for LUAD (reviewed in(Rowbotham and Kim, 2014), restricting Cre recombinase expression to either club cells (CL:0000158) or pulmonary alveolar type II (AT2; CL:0002063) cells in KP mice results in LUAD, but in different locations and with distinct phenotypic characteristics (Sutherland et al., 2014). The median survival of KP mice following expression of Cre in club cells or AT2 cells is 25.3 and 23.1 weeks, respectively. KP mice with expression of Cre recombinase restricted to club cells developed metastases in the myocardium and the thoracic cavity at an unspecified frequency at 25.2 weeks after tumor induction (Jackson et al., 2005). In humans, the four most common sites of LUAD metastasis are the lymph nodes, brain, bone, and liver (Guan et al., 2025, Liu et al., 2024).

Tumor progression requires intercellular communication: both local, between the primary tumor and the tumor microenvironment (TME), and long-distance, between the tumor and sites of future metastases (reviewed in (Wortzel et al., 2019, Sole and Lawrie, 2021). MicroRNAs (miRNAs) are small RNAs that regulate multiple cancer-relevant processes, including gene expression, cell invasion, proliferation, migration, cell viability, epithelial-to-mesenchymal transition (EMT), metastasis, and therapeutic resistance (reviewed in (El Founini et al., 2021, Zhang et al., 2024, Durendez-Saez et al., 2021). MiRNAs can be transported intercellularly in extracellular vesicles (EVs), transferred to recipient cells, and alter gene expression within recipient cells (Valadi et al., 2007) (reviewed in(O’Brien et al., 2022). Although miRNAs have been implicated in the progression and metastasis of LUAD (reviewed in (Martinez-Espinosa et al., 2023), the contributions of individual miRNAs and miRNA-regulated pathways are largely unknown. To address this knowledge gap, we investigated the contribution of global miRNA disruption to LUAD progression.

Biogenesis of mature, functional miRNAs relies upon the activity of DICER1 (HGNC:17098), an RNase III enzyme. Germline mutations in human *DICER1* are associated with a variety of rare benign and malignant neoplasms, collectively referred to as DICER1 tumor-predisposition syndrome. These tumors frequently harbor a second, somatic variant within the RNase IIIb domain of *DICER1,* which alters the ratio of 3p and 5p miRNAs expressed in affected tissues (reviewed (Kim et al., 2017, Jansen et al., 2025). Heterozygous germline loss-of-function (LOF) variants in the *DICER1* gene characterize pleuropulmonary blastoma (PPB; MIM:601200), a rare pediatric lung tumor (Klein et al., 2014, Caroleo et al., 2020, Gonzalez et al., 2022) (reviewed in (Foulkes et al., 2014, Gonzalez et al., 2022). PPB germline *DICER1* mutations occur in mesenchymal tumor cells, while the somatic *DICER1* mutations are found within the tumor-associated epithelial cells, suggesting a non-cell-autonomous mechanism of cancer initiation (Hill et al., 2009). PPB arises during fetal lung development and can temporally evolve from a type I multicystic lesion to a type III malignant solid tumor (Hill et al., 2009) (reviewed in (Gonzalez et al., 2022). Biallelic *DICER1* mutations appear to be insufficient for progression of type I PPB tumors to types II or III, suggesting that disrupting DICER1 function works in concert with additional mutations to facilitate lung tumor progression (Gonzalez et al., 2022). Sequencing of mesenchymal tumor tissues from PPB patients has identified mutations in *TP53* and *NRAS* as the most frequent additional drivers of PPB progression (Pugh et al., 2014).

Here, we describe novel genetically engineered mouse models of *Kras*-driven LUAD which exhibit differences in tumor progression rates and expected length of survival. The models are derived from a parental mouse strain carrying conditional alleles of *Kras, Trp53*, and *Dicer1* (KPD strain) on a C57BL/6J background. Following Cre exposure, KPD mice express one allele of oncogenic *Kras^G12D^*, lack functional *Trp53*, and retain only one functional *Dicer1* allele. We mutated the remaining *Dicer1* allele in exon 24, one of two exons harboring hotspot mutations in PPB. Using genetic Cre drivers and adenoviral and lentiviral vectors, we restricted Cre expression and *Dicer1* mutation to specific cell types in the lungs. We observed accelerated tumor progression and decreased expected survival only when we expressed Cre in club cells and mutated *Dicer1* in AT2 cells. Induction of tumorigenesis in AT2 cells with *Dicer1* truncation in club cells modestly accelerated tumor progression, without affecting expected survival. These mouse models represent an important new resource for investigating the cell-type-dependent and non-cell-autonomous contributions of *Dicer1* mutation, and subsequent miRNA disruption, to lung cancer progression.

## RESULTS

### Lentiviral vectors efficiently mutate *Dicer1* in exon 24

We designed a CRISPR guide RNA, x24sgRNA1, to target the region of murine *Dicer1* exon 24 (Fig. 1A) that is homologous to the region of human *DICER1* that is mutated in PPB patients. Our choice of guide RNA was not intended to create a mouse model of PPB, but to induce a *Dicer1* truncation known to be relevant to lung cancer. We produced a total of four lentiviral vector constructs (Fig. 1B): (1) pLex307iCre, which ubiquitously expresses Cre from the EF-1a promoter but does not truncate *Dicer1*; (2) pLCx24Cas9, which ubiquitously expresses Cre and Cas9 from an EFS promoter as well as x24sgRNA1 from a human U6 (hU6) promoter in infected cells; (3) SPCx24, which expresses x24sgRNA1 from an hU6 promoter and *Cas9* from an SPC promoter in infected AT2 cells; and (4) CC10×24, which expresses x24sgRNA1 from an hU6 promoter and Cas9 from aCC10 promoter in infected club cells.

**Fig. 1.**
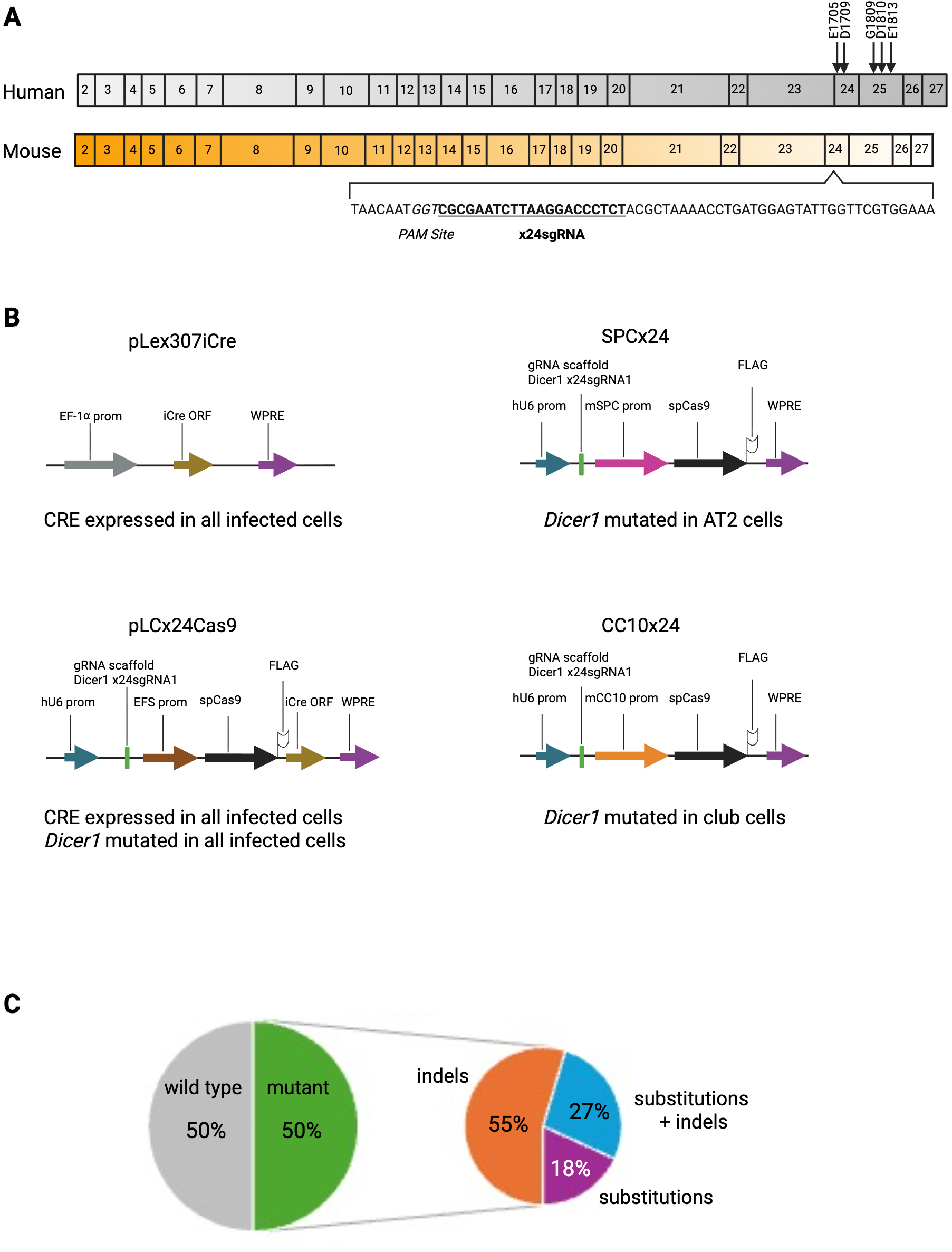
Dicer1 domains, truncating viral vectors and sequencing results. (A) Graphical representation of coding exons in human (top) and mouse (bottom) DICER1. Arrows show the approximate locations of the five human somatic hotspot mutations (PMID 33293352). Exons 24 and 25 of both human and mouse DICER1 encode the RNase IIIb domain (PMID 33293352). The location and sequence of the x24sgRNA (underlined bold text) is also shown relative to the Protospacer Adjacent Motif (PAM) site (italics) (B) Important functional elements of the four viral vectors generated for this study: ORF, open reading frame; WPRE, Woodchuck Hepatitis Virus Post-transcriptional Regulatory Element; prom, promoter; FLAG, FLAG-tag. The cell types which express Cre recombinase or mutated DICER1 after infection with each viral vector are also indicated. (C) Results of cas9 capture sequencing shows that *Dicer1* is mutated in one half of all reads covering the targeted region. Sequencing reads containing mutations are further broken down into insertions and deletions (indels), nucleotide substitutions or both substitutions and indels.

To test the efficiency of our CRISPR guide RNA, we generated the Kras^tm4Tyj/+^;Trp53^tm1Brn/tm1Brn^;Dicer1^tm1Bdh/+^;Gt(ROSA)^26Sortm4(ACTB-tdTomato,-EGFP)Luo/+^ strain (KPDT; Table 1). KPDT mice carry the dual-fluorescent Cre reporter allele mT/mG, which expresses the red fluorescent protein tdTomato in every cell (Muzumdar et al., 2007). Upon exposure to Cre recombinase, cells turn off expression of tdTomato and turn on expression of enhanced green fluorescent protein (GFP), marking Cre-expressing cells. Cre expressing cells are the cell population which express *Kras^G12D^*, delete *Trp53* and, thus, form tumors. After infecting a KPDT mouse with Ad5mSPC-Cre and CC10×24, we made a single lung cell suspension and used fluorescence-activated cell sorting (FACS) to separate cells into GFP^+^ tumor cell and tdTomato^+^ non-tumor cell fractions. An antibody against Epithelial Cell Adhesion Molecule (EpCAM) was used to further separate the tdTomato^+^ cell population into epithelial (EpCAM high) and non-epithelial (EpCAM low) populations.

**Table 1.** Summary of mouse models used in this study and their cell type specific alterations.

| Model<br>Name | Mouse<br>Strain | Cre<br>Viral Vector | *Dicer1*<br>Viral Vector | Tamoxifen? | Cell Type |  |  |  |
| --- | --- | --- | --- | --- | --- | --- | --- | --- |
|  |  |  |  |  | KRAS<sup>G12D</sup> | TRP53<br>del | DICER1<br>del | DICER1<br>Mut |
| KP (adeno) | KP | Ad5CMVCre | none | no | various | various | none | none |
| KP (lenti) | KP | pLex307iCre | none | no | various | various | none | none |
| KPD | KPD | pLex307iCre | none | no | various | various | various | none |
| KPD<sup>del/mut</sup> | KPD | pLCx24Cas9 | pLCx24Cas9 | no | various | various | various | various |
| KPDS | KPDS | none | SPCx24 | yes | club | club | club | AT2 |
| KPDSCre | KPDS | none | none | yes | club | club | club | none |
| KPDSDicer1 | KPDS | none | SPCx24 | no | none | none | none | AT2 |
| KPDF | KPDF | none | CC10x24 | yes | AT2 | AT2 | AT2 | club |
| KPDFControl | KPDF | none | none | no | none | none | none | none |
| KPDT-1 | KPDT | Ad5CC10-Cre | SPCx24 | no | club | club | club | AT2 |
| KPDT-1Cre | KPDT | Ad5CC10-Cre | none | no | club | club | club | none |
| KPDT-1Empty | KPDT | Ad5CMVEmpty | none | no | none | none | none | none |
| KPDT-2 | KPDT | Ad5SPC-Cre | CC10x24 | no | AT2 | AT2 | AT2 | club |
| KPDT-2Cre | KPDT | Ad5SPC-Cre | none | no | AT2 | AT2 | AT2 | none |
| KPDT-2Empty | KPDT | Ad5CMVEmpty | none | no | none | none | none | none |
| C57BL/6J | C57BL/6J | none | none | no | none | none | none | AT2 |

*Cas9*-targeted sequencing of approximately 1.4 million EpCAM-high, tdTomato^+^ cells identified 22 reads that covered the binding site for x24sgRNA1 (Lesbirel et al., 2021). Of these reads, 11 (50%) contained mutations within ten nucleotides of the predicted cut site for x24sgRNA1(Fig. 1D). These mutation-containing reads were categorized as insertions and deletions (indels; 6/11), substitutions (3/11), or substitutions plus deletions (2/11). Under these experimental conditions, our CC10×24 lentiviral vector mutated *Dicer1* in half of the tdTomato^+^ EpCAM-high cell population, which likely contained cell types in addition to club cells. Therefore, we conclude that our x24sgRNA1 CRISPR guide RNA efficiently induces mutations in exon 24 of *Dicer1* as designed.

### Alteration of *Dicer1* in lung tumor cells has a modest effect on pulmonary tumorigenesis

In the previously characterized KP mouse strain, intranasal instillation of an adenoviral vector expressing Cre under the ubiquitous cytomegalovirus promoter (Ad5CMVCre; University of Iowa) resulted in all animals developing multiple tumors, ranging from well-differentiated to highly dysplastic lesions (Jackson et al., 2005). The authors noted lymph node metastases in over 50% of infected animals, as well as a small percentage of animals with metastases to distant organs, including the kidneys. We attempted to repeat these reported results by infecting 5 male and 5 female KP mice with 5 × 10^6 plaque-forming units (pfu) of Ad5CMVCre at 7 weeks of age (cohort 1; Table 2). For clarity, we refer to KP strain mice intranasally infected with Ad5CMVCre as the KP (adeno) mouse model (Table 1). One KP (adeno) female was found dead with lung lesions at 12 weeks after viral infection. At 16 weeks after viral infection, 8 of 9 animals had pulmonary lesions consisting of a mixture of adenomas and adenocarcinomas, with regions of hyperplasia, in good agreement with previously reported results (Fig. 2A, Table 2). However, we were unable to replicate the published high metastasis rate (>50% of infected animals) in the KP (adeno) model. We examined heart and thymus tissues from nine KP (adeno) animals and thoracic lymph nodes from six animals and observed only one metastatic lesion outside of the lungs, which we found in a lymph node (Fig. 2B). Genetic modifiers may partially explain this discrepancy in the frequency of metastasis as Jackson et al. generated their KP mice on a mixed 129S/C57BL/6J background while our KP (adeno) and subsequent models are on a fixed C57BL/6J background. It is also possible that importation and rederivation at JAX altered the lung microbiota of these mouse strains. Commensal microbiota within the lungs of KP mice have been shown to drive local inflammation and lung tumor progression (Jin et al., 2019).

**Table 2.** Summary of mouse cohorts used in this study.

| Cohort # | Mouse Model | # of Animals | Age at Harvest | Animals With Tumors | Lymph Node Mets | Died Before Harvest |
| --- | --- | --- | --- | --- | --- | --- |
| 1 | KP (adeno) | 5M/5F | 16 weeks | 8/9 | 1/6 | 1 F |
| 2 | KP (lenti) | 5M/8F | 16 weeks | 6/13 | 0/5 |  |
| 3 | KPD | 4M/6F | 16 weeks | 3/10 | 0/3 |  |
| 4 | KPD<sup>del/mut</sup> | 10M/5F | 16 weeks | 5/15 | 0/1 |  |
| 5 | KPDS | 7M/6F | 12 weeks | 12/12 | 4/7 | 1F/2M |
| 6 | KPDS | 6M/5F | 10 weeks | 11/11 | 1/2 |  |
| 7 | KPDSCre | 7M/2F | 10 weeks | 8/9 | 1/2 |  |
| 8 | KPDSDicer1 | 3M/10F | 16 weeks | 9/13 | 0/1 |  |
| 9 | C57BL6/J | 5M/4F | 16 weeks | 0/9 | 0/2 |  |
| 10 | KPDFControl | 7M/7F | 2-8 weeks | 9/11 | 0/0 | 2F/1M |
| 11 | KPDT-1 | 5M/5F | 10 weeks | 9/10 | 0/3 |  |
| 12 | KPDT-1Cre | 5M/5F | 10 weeks | 0/9 | 0/8 |  |
| 13 | KPDT-1Empty | 5M/5F | 10 weeks | 0/10 | 0/1 |  |
| 14 | KPDT-2 | 5M/5F | 20 weeks | 9/10 | 0/9 |  |
| 15 | KPDT-2Cre | 5M/5F | 20 weeks | 8/9 | 0/7 | 1 M |
| 16 | KPDT-2Empty | 5M/5F | 20 weeks | 0/10 | 0/0 |  |
| 17 | KPDT-2 | 3M/5F | 10 weeks | 2/8 | 0/3 |  |
| 18 | KPDT-1 | 2M/1F | 11 weeks | 3/3 | 0/3 |  |
| 19 | KPDT-1 | 2M/1F | 12 weeks | 3/3 | 0/3 |  |

**Fig. 2.**
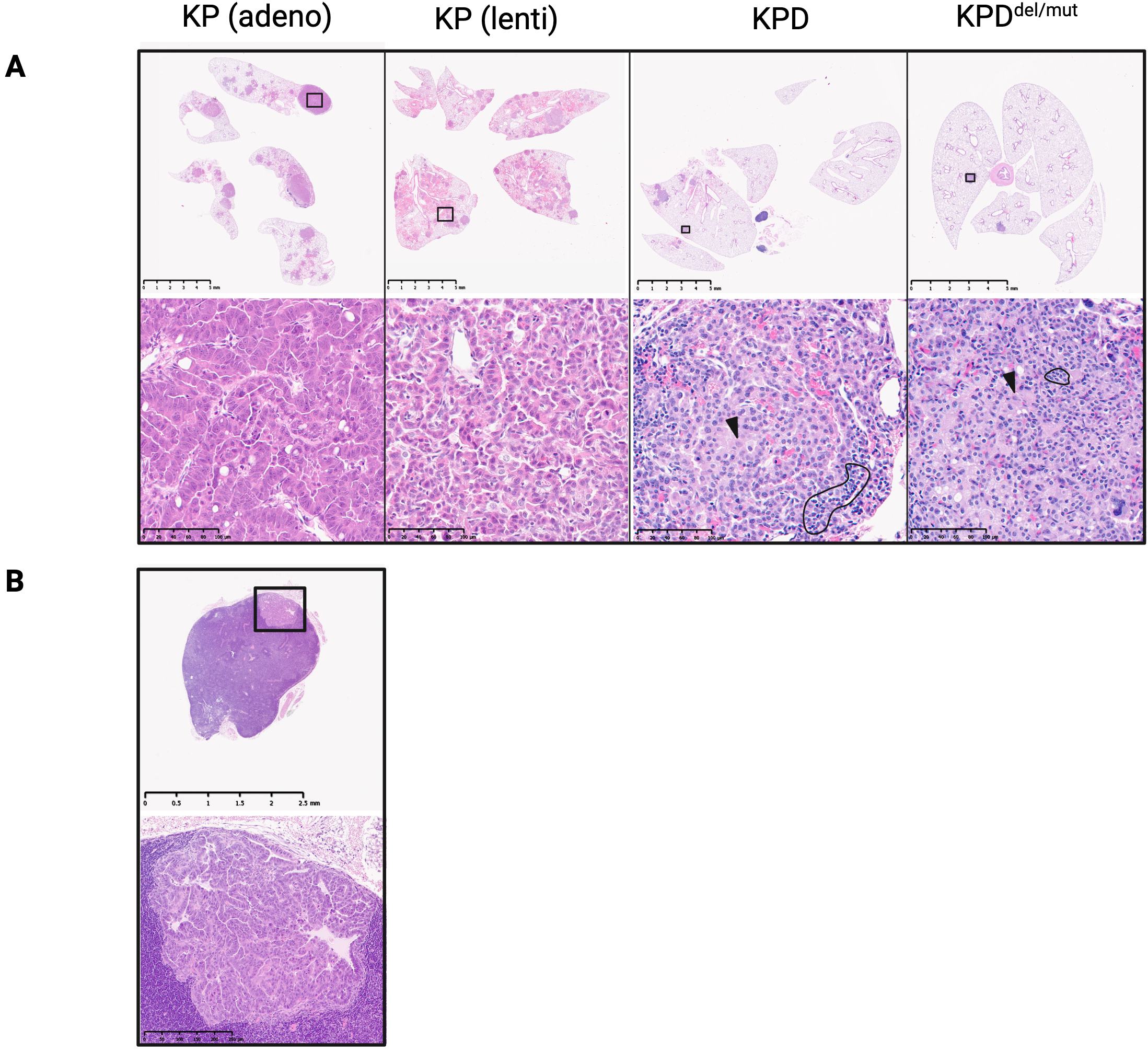
Histological comparison of tissues from four di;erent mouse models of pulmonary adenocarcinoma. For all four models, mice were infected at 7-8 weeks of age and tissues were collected at 16 weeks after infection, sectioned and stained with H&E. (A) Lung tissues from KP (adeno) mice showed a mix of adenomas and adenocarcinomas with regions of hyperplasia that are shown at lower magnification (top row) and boxed areas at higher magnification (bottom row). Arrowheads indicate representative tumor cells and examples of infiltrating inflammatory cells in lung tissues from KPD and KPD^del/mut^ mice are circled (black lines). (B) Lymph node tissue from the only mouse in all four cohorts in which a metastasis (boxed region) was detected is shown at lower magnification (top row) and at higher magnification (bottom row). Lung and lymph node tissues shown in panels A and B are from the same KP (adeno) mouse.

Since our viral vectors for mutating *Dicer1* (Fig. 1B) are lentiviruses, we generated a lentiviral vector, pLex307iCre (Fig. 1B), that expresses Cre recombinase in a variety of infected cell types to more accurately compare our different cohorts of mice. We infected 13 KP mice, 8 females and 5 males, with pLex307iCre at 7 weeks of age and harvested them at 16 weeks post-viral infection (cohort 2). We refer to KP mice intranasally infected with pLex307iCre as the KP (lenti) mouse model (Table 1). Histological examination of the lungs of KP (lenti) animals determined that one mouse had focal hyperplasia. Six additional animals had tumors, which were a mix of adenomas and adenocarcinomas, some with large, multinucleated cells (Fig. 2A, Table 2). Comparison of the lung tissues from mice of cohorts 1 and 2 suggests that our lentiviral vector is less efficient at inducing lung tumors than Ad5CMVCre, but the tumors initiated by both viral vectors are histologically comparable. We examined the thymus and hearts from all cohort 2 animals plus the thoracic lymph nodes from five animals and did not observe evidence of metastasis outside the lungs.

*Dicer1* has been described as a haploinsufficient tumor suppressor because monoallelic, but not biallelic, *Dicer1* loss promotes tumorigenesis in a non-cell-autonomous fashion (Kumar et al., 2009, Yoshikawa et al., 2013, Lambertz et al., 2010). To examine the effects of ablating one allele of *Dicer1* on pulmonary tumorigenesis, we added one floxed allele of *Dicer1* to KP mice to generate the *Kras^tm4Tyj/+^*;*Trp53^tm1Brn/tm1Brn^;Dicer1^tm1Bdh/+^*mouse strain (KPD). We intranasally infected 10 KPD^+/Fl^ animals (6 females and 4 males) with pLex307iCre at 7 weeks of age and harvested them at 16 weeks post-infection (cohort 3). We refer to KPD animals intranasally infected with pLex307iCre as the KPD mouse model (Table 1). Conditional deletion of one allele of *Dicer1* in pulmonary tumor cells resulted in fewer animals (30% vs. 46%) having tumors, not more, and each animal had visibly fewer tumors than animals in cohort 2 (Fig. 2A, Table 2). Our results agree with a recently reported murine model of fusion-negative rhabdomyosarcoma (FN_RMS), which showed that germline heterozygous loss of *Dicer1* promoted tumor formation, whereas conditional heterozygous deletion within tumor cells did not (Larsen et al., 2025). Thus, a single heterozygous LOF mutation in *Dicer1* within tumor cells has a modest effect on tumorigenesis.

Tumors in cohort 3 animals were a mix of adenomas and carcinomas with large multinucleated cells. In comparison to tumors in KP (adeno) and KP (lenti) mice, tumors in KPD mice had more infiltrating inflammatory cells, including lymphocytes, plasma cells, and mononuclear cells (Fig. 2A). The increased presence of inflammatory cells was consistent across all tumor-bearing KPD animals and across all tumors within the same animal. Despite evidence of tumor cells invading the airways, we did not observe metastasis to the lymph nodes.

Next, we tested the effects of mutating the remaining allele of *Dicer1* within tumor cells of KPD mice by infecting them with our pLCx24Cas9 construct to generate a mouse model that we refer to as KPD^del/mut^ (Table 1). In KPD^del/mut^ animals, tumors express *Kras^G12D^*, delete both Trp53 alleles and one Dicer1 allele, and express *Dicer1* from the remaining mutated allele. Histological examination of 15 KPD^del/mut^ animals, 5 females and 10 males, harvested at 16 weeks after viral infection (cohort 4), revealed that 2/15 animals had focal hyperplasia, and an additional 5 animals had tumors (Table 2). Tumors were a mix of adenomas and adenocarcinomas containing potentially multinucleated cells with irregular borders (Fig. 2A). Again, we saw an increased presence of inflammatory cells in these mice compared to KP (adeno) and KP (lenti) mice with wild-type DICER1 expression (Fig. 2A). We also observed tumor cells invading the airways of KPD^del/mut^ animals, but we did not detect metastasis outside of the lungs.

### Initiating tumors in Club cells and mutating *Dicer1* in ATII cells shortens survival and accelerates tumorigenesis

To determine if *Dicer1* mutation affects tumor progression in a non-cell autonomous fashion, potentially by altering intercellular communication, we induced pulmonary tumors in club cells and mutated *Dicer1* in ATII cells. To restrict expression of Cre to club cells, we mated KPD mice to mice that express a tamoxifen-inducible form of Cre from the Club cell specific Secretoglobin1a1 (*Scgb1a1*) locus to generate the *Kras^tm4Tyj/+^;Trp53^tm1Brn/tm1Brn^;Dicer1^tm1Bdh/+^;Scgb1a1^tm1(cre/ERT)Blh/+^*strain, (KPDS; Table 1). We administered four doses of tamoxifen by oral gavage to 13 KPDS animals, 6 females and 7 males, at seven weeks of age then intranasally infected them with 2 x 10^4^ TU of SPCx24 the following week (cohort 5), which we refer to as the KPDS mouse model (Table 1). At 12 weeks after the first tamoxifen dose, KPDS animals developed respiratory distress. We were unable to recover organs from one animal found dead and were able to recover only the lungs from two additional animals that were found dead (Table 2). In all necropsied KPDS animals, approximately 40-50% of lung tissue was comprised of predominantly adenocarcinomas, some with multinucleated giant cells, plus a few adenomas and we observed tumor cells invading and occluding the airways (Fig. 3A). Histological analysis of the thoracic lymph nodes identified metastases in four of seven KPDS animals (Fig. 3B). Many animals also had areas of cellular atypia, but no metastases, within their livers (data not shown). We did not observe metastasis within the heart, thymus, kidneys, adrenals or spleen tissues in any animal of this cohort (data not shown). We were able to reproduce the observed decrease in survival and presence of lymph node metastases when we reduced the number of tamoxifen doses from four to two (cohort 6, data not shown).

**Fig. 3.**
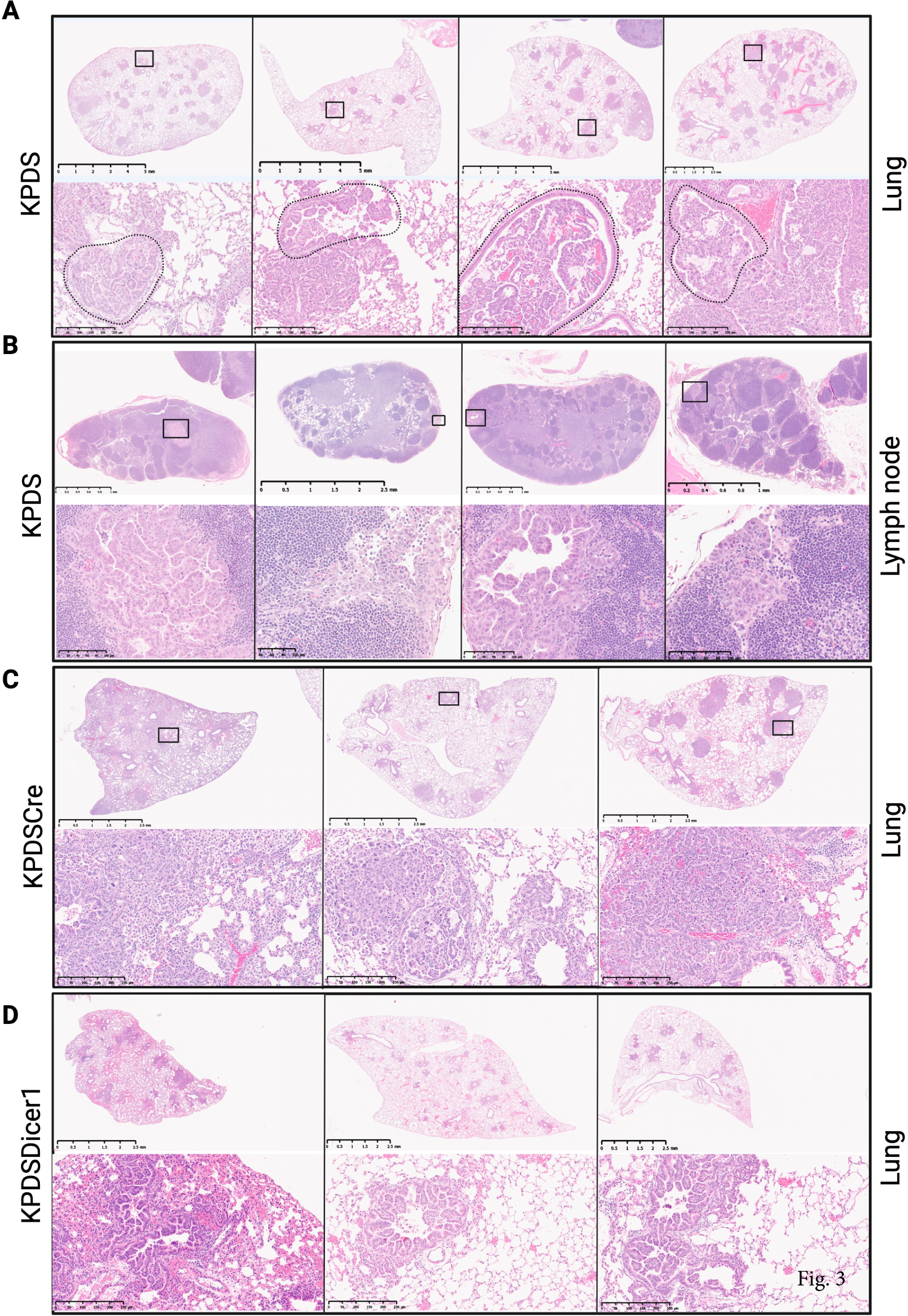
Histological comparison of tissues from KPDS, KPDS Cre and KPDS Dicer1 mouse models. Harvested lung tissues were embedded, sectioned and stained with H&E. Top rows of each panel show tissues at a lower magnification and boxed areas are shown at a higher magnification in lower rows. (A) KPDS mice were administered tamoxifen at 7 weeks of age, SPCx24 the following week and harvested at 12 weeks after the first dose of tamoxifen. Lung tissues from four KPDS mice (vertical columns) show predominantly adenocarcinomas, some with multinucleated giant cells (top panel). Tumor cells are invading and filling up airways (dotted lines, lower row). (B) Lymph nodes from 57% of KPDS mice harbored metastases (boxed areas, upper row) which are shown at higher magnification (lower row). Tissues from the same mice are shown in both panels A and B. (C) KPDSCre mice were administered tamoxifen at 7-8 weeks of age and harvested at 10 weeks after the first dose of tamoxifen. Lung tissues from three KPDS Cre mice display predominantly adenocarcinomas, shown at lower (upper row) and higher (lower row) magnification. (D) KPDSDicer1 were administered SPCx24 at 7-8 weeks of age KPDSCre mice were harvested at 16 weeks after receiving virus. Lung tissues from three KPDSDicer1 animals with adenomas and a few early-stage carcinomas (upper row). Higher magnification (lower row) shows variation in size of nuclei and cells but no mitotic figures.

As controls, KPDS animals received either tamoxifen (KPDSCre), or SPCx24 alone (KPDSDicer1) (Table 1). Administration of four doses of tamoxifen resulted in 8 of 9 KPDSCre animals with pulmonary tumors and one animal with a lymph node metastasis (cohort 7, Table 2, Fig. 3C). Tumors from animals in cohorts 3 and 7 were histologically identical, indicating that restricting tumor initiation to club cells does not alter tumor histology. Surprisingly, mutating *Dicer1* within ATII cells of KPDS animals, in the absence of inducing Cre expression, resulted in 9 of 13 KPDSDicer1 animals with abnormal lung histology, including areas of diffuse hyperplasia, adenomas, and early-stage adenocarcinomas (cohort 8, Fig. 3D). However, mutating *Dicer1* within ATII cells of C57BL/6J animals did not result in tumors (data not shown), suggesting that expression of the *Scgb1a1* Cre driver in the various KPDS models was leaky. Analysis of lung tissues from cohort 8 animals confirmed the presence of a small amount of recombined *Kras* (Fig. S1) in the absence of tamoxifen administration.

To test if reversing the cell types that harbor *Dicer1* mutations or bear tumors would also affect tumor progression, we restricted expression of Cre to ATII cells by breeding KPD mice to mice that express a tamoxifen inducible form of Cre from the ATII cell specific Surfactant Protein C (*Sftpc*) locus to generate the *Kras^tm4Tyj/+^;Trp53^tm1Brn/tm1Brn^;Dicer1^tm1Bdh/+^;Sftpc1^tm1(cre/ERT2)Blh/+^* strain (KPDF; Table 1). We intended to administer four doses of tamoxifen by oral gavage to KPDF animals at seven weeks of age, then intranasally infect them with 2 x 10^4^ TU of CC10×24 the following week to generate a model also referred to as KPDF. However, the *Sftpc* Cre driver proved so leaky that the majority (9/11) of KPDF animals administered only corn oil at 7 weeks of age, referred to as the KPDFControl model, developed adenocarcinomas and began dying by 9 weeks of age. PCR analysis of tumors from these mice confirmed the presence of both recombined *Kras* (Fig. S1) and recombined *Trp53* in the absence of tamoxifen administration (data not shown). As in KPDS mice, the recombination we observed in KPDFControl animals varied between individual mice.

### Lung tumor progression is accelerated only when tumors are initiated in club cells and *Dicer1* is mutated in ATII cells

Given the leakiness of the *Sftpc* and *Scgb1a1* Cre drivers (described above), we switched to a dual viral vector approach to induce Cre expression and mutate *Dicer1* in different lung cell populations. These mouse models used the KPDT mouse strain, which we previously generated for testing the efficiency of our CRISPR guide RNA. For the KPDT-1 model, we expressed Cre in club cells and mutated *Dicer1* in ATII cells by infecting KPDT mice with Ad5CC10-Cre (University of Iowa) and SPCx24. In the KPDT-2 model, we reversed the cell types by infecting KPDT mice with Ad5mSPC-Cre and CC10×24 (Table 2). In mice, club cells reside within the proximal region of the respiratory system, while ATII cells reside in the distal, alveolar region.

The effects of *Dicer1* mutation on tumor progression are cell-type specific. Previously, Sutherland et al. determined that the median survival for KP mice infected with either Ad5mCC10-Cre or Ad5SPC-Cre was 25.3 and 23.1 weeks, respectively (Sutherland et al., 2014). Deletion of one allele of *Dicer1* in tumor cells initiated in club cells combined with mutation of *Dicer1* in ATII cells (cohort 11) resulted in 90% of animals (9/10) having a mix of predominantly adenocarcinomas plus lesser amounts of adenomas and a few areas of hyperplasia by 10 weeks after viral infection (Fig. 4A). Several of the carcinomas appeared aggressive. These tumors often surrounded and invaded airways, sometimes presenting as “rafts” of tumor cells within the airways (Fig. 4A, left panel). In alignment with our animal use protocol, tumor-bearing mice were not allowed to reach the moribund stage. However, to better determine how long KPDT-1 mice might survive after tumor induction, we euthanized animals from two small additional cohorts, at 11 (cohort 18, data not shown) and 12 weeks after viral infection (cohort 19, Fig. 4B), respectively. By 11 weeks after tumor induction, animals displayed very aggressive carcinomas, regardless of tumor size, that were invading blood vessels. Two animals, one each from cohorts 18 and19, displayed large areas of hemorrhage and blood clots that occluded blood vessels. One mouse from cohort 19 went into respiratory distress on the day of euthanasia, suggesting that 12 weeks after viral infection is reaching the maximum survival time for KPDT-1 animals. We also observed tertiary lymphoid structures near the lungs in several animals from cohorts 18 and 19. In contrast, all animals infected only with Ad5mCC10-Cre and harvested 10 weeks later (cohort 12) did not have tumors, while one animal had a region of hyperplasia (Fig. 4C). Thus, deletion of one allele of *Dicer1* in tumor cells initiated in club cells combined with mutation of *Dicer1* in ATII cells accelerated tumor progression and reduced expected survival compared to initiating tumors in club cells alone. As expected, no abnormal pathology was observed in animals infected with Ad5Empty viral vector (cohort 13, Fig. 4D).

**Fig. 4.**
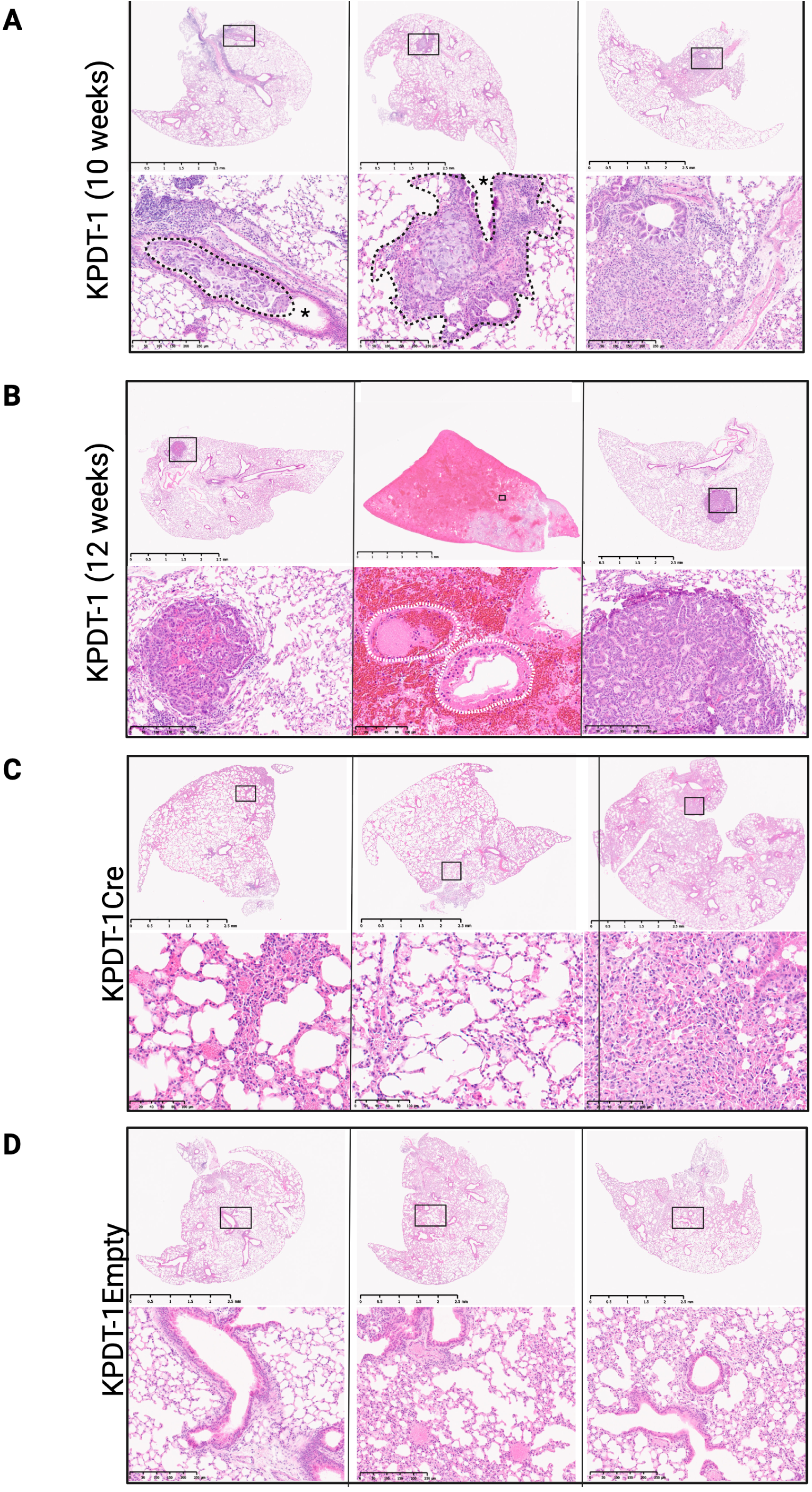
Dicer1 mutations in ATII cells accelerates progression of tumors initiated in Club cells. Lung tissues are shown from three mice each (vertical rows) from four dieerent models (horizontal panels), all infected at 6-8 weeks of age and harvested at 10 (panels A, C and D) or 12 weeks (panel B) after viral infection. Boxed regions from upper rows of each panel (lower magnification) are shown at higher magnification in lower rows. (A) Tumors in lungs from KPDT-1 animals, in which tumors are initiated in Club cells and Dicer1 is mutated in ATII cells, were a mix of adenomas and carcinomas, with tumors invading (dotted black line lower left panel) or surrounding (dotted black line lower middle panel) airways (asterisk). (B) At 12 weeks after viral infection, tumors are invading blood vessels (dotted white line middle panel) and some lung lobes display hemorrhage (middle panel). (C) Lung tissues from KPDT-1Cre, in which tumors are initiated in Club cells, but Dicer1 is wild type, showed no tumors. (D) Lung tissues from KPDT-1Empty animals, without tumor initiation or mutated Dicer1, show no abnormal pathology.

When we reversed cell types harboring *Dicer1* mutations or bearing tumors in the KPDT-2 model, *Dicer1* mutation had modest effects on tumor progression and expected survival. At 20 weeks after infection, 90% of KPDT-2 animals (9/10) had carcinomas (cohort 14, Fig. 5A). While some of these tumors were quite large, we did not observe them growing around or into airways, nor did we ever observe tumors invading blood vessels. We harvested KPDT-1 animals at 10 weeks after viral infection because they appeared unable to survive for 20 weeks. Therefore, we collected an additional cohort of KPDT-2 animals (cohort 17), consisting of 5 females and 3 males, at 10 weeks after viral infection (Fig. 5B) so that we could compare tumor development in KPDT-1 and KPDT-2 animals at the same time point after viral infection. At 10 weeks after viral infection, 3 of 8 KPDT-2 animals developed tumors, mostly adenomas with a few small adenocarcinomas. There were also areas of hyperplasia in several animals. Thus, tumors in KPDT-1 animals are more aggressive than tumors in KPDT-2 animals at 10 weeks after viral infection. We observed a similar percentage of animals with tumors (8/9) at 20 weeks after infection with only Ad5SPC-Cre (cohort 15, Fig. 5C). One additional animal from cohort 15 was found dead with lung tumors the day before scheduled tissue harvest. Mutation of *Dicer1* in club cells appears to have a small effect on progression of tumors initiated in ATII cells, as tumors in animals from cohort 14 were slightly more aggressive than those in animals from cohort 15, but no noticeable effect on expected survival as KPDT-2 animals did not show signs of respiratory distress at 20 weeks after viral infection compared to the previously reported survival of 23.1 weeks. No abnormal histology was observed in animals infected with Ad5Empty (cohort 16, Fig. 5D).

**Fig. 5.**
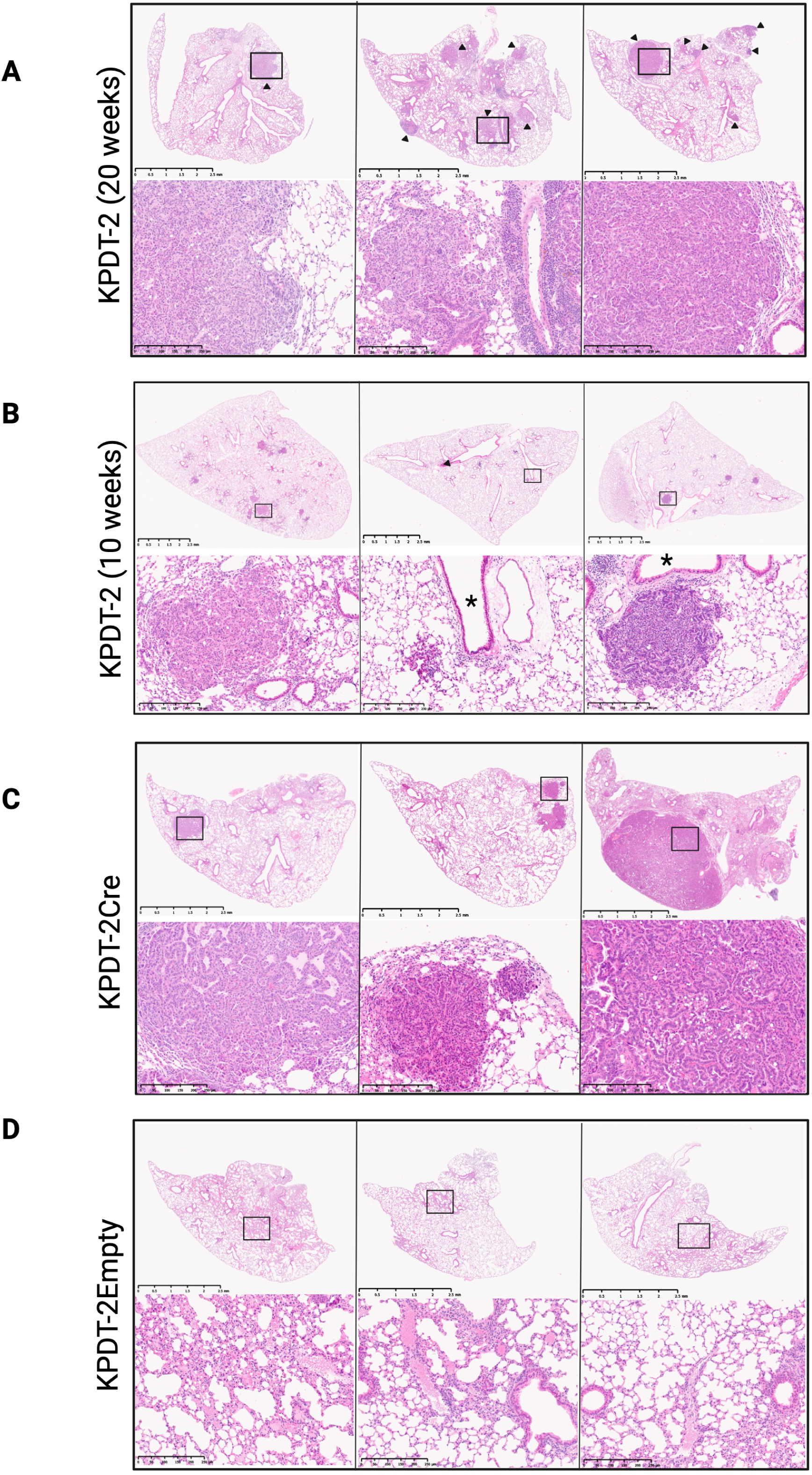
Dicer1 mutations in Club cells have a modest e;ect on progression of tumors initiated in ATII cells. Lung tissues are shown from three mice each from three dieerent models, all infected at 6-8 weeks of age and harvested at either 10 (panel B) or 20 weeks (panels A, C and D) after infection. Boxed regions from upper rows of each panel (lower magnification) are shown at higher magnification in lower rows. (A) Lungs from KPDT-2 animals, in which tumors are initiated in ATII cells and Dicer1 is mutated in Club cells, collected at 20 weeks after infection shows predominantly adenocarcinomas (arrow heads, upper rows). (B) At 10 weeks after tumor induction, lungs of KPDT-2 animals have a mixture of areas of hyperplasia, adenomas and a few carcinomas, none of which are invading the airways (asterisk). (C) Lung tumors in KPDT-2Cre animals, in which tumors are initiated in ATII cells, are predominantly adenocarcinomas by 20 weeks after tumor induction. (D) Lung tissues from KPDT-2Empty animals, without tumor initiation or mutated Dicer1, show no abnormal pathology.

## DISCUSSION

Here we describe a series of *Kras*-driven mouse models of pulmonary adenocarcinoma that we generated to investigate the effects of disrupting miRNA expression within different cell types on tumor progression. Due to leaky Cre expression, we abandoned models using the *Scgb1a1* and *Sftpc* tamoxifen-inducible Cre drivers and developed a dual-virus-mediated engineering strategy to generate our KPDT-1 and KPDT-2 models. Our findings demonstrate that mutation of *Dicer1* accelerates tumor progression and reduces expected survival in KP mice in a cell-type-dependent manner (Fig. 6). Tumor progression and expected survival were affected only when mice harbored two distinct *Dicer1* alterations, each within different cell populations. In contrast to previously described mouse tumor models, our KPDT-1 and KPDT-2 models harbor a cell-type-specific loss-of-function (LOF) mutation in one *Dicer1* allele in one cell type and an RNase III domain mutation in *Dicer1* in a second cell type. Thus, both *Dicer1* LOF and RNase III domain mutations affect tumorigenesis, and their effects are cell-type-dependent. Our results support the hypothesis that *Dicer1* alteration either enhances miRNA-mediated intercellular communication required for tumor progression or disrupts communication necessary for tumor inhibition.

**Fig. 6.**
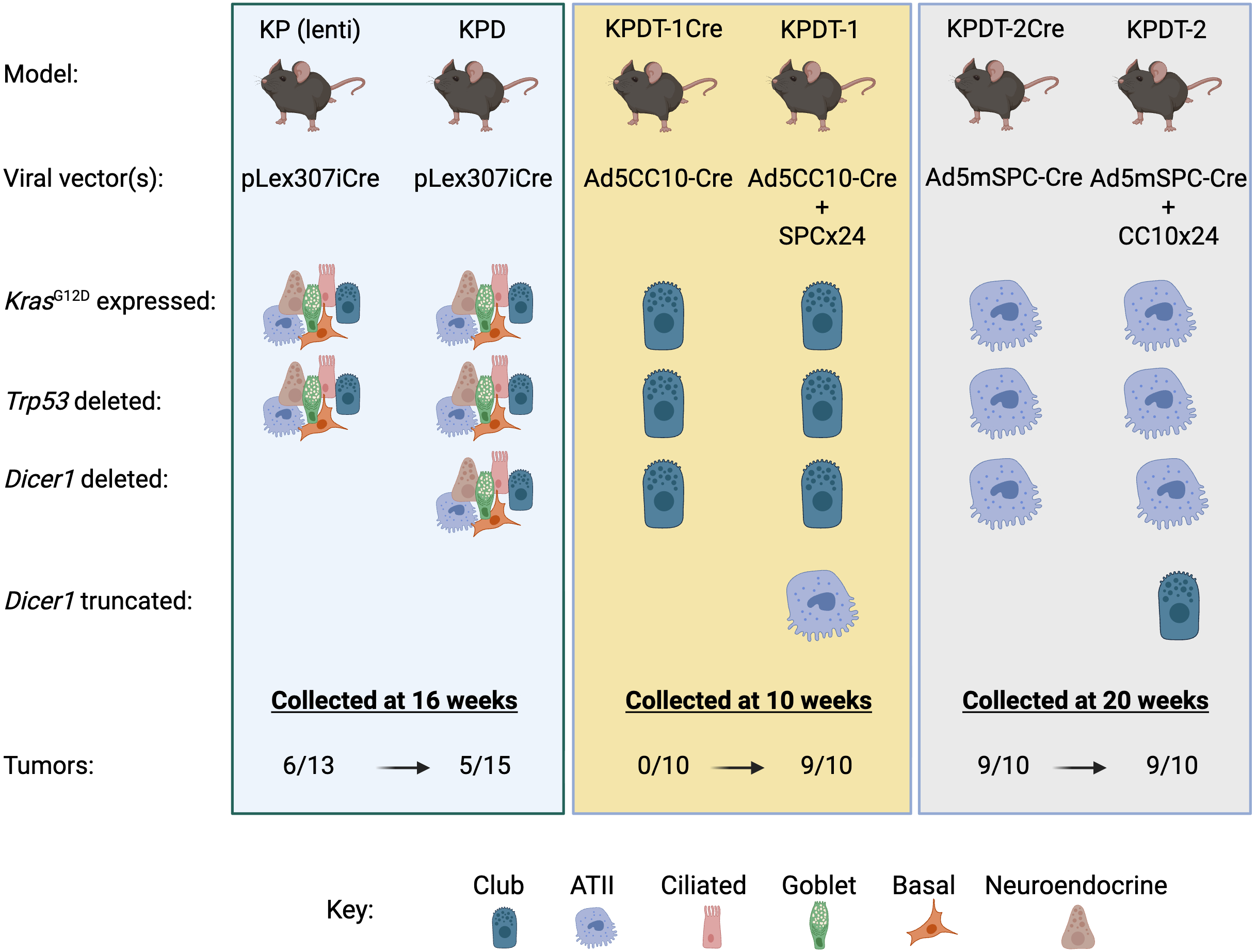
Summary of KP, KPD, KPDT-1 and KPDT-2 mouse models of pulmonary adenocarcinoma. Our mouse models of pulmonary adenocarcinoma are based upon the KP model, in which Cre controls expression of oncogenic *Kras*^G12D^ and deletion of *Trp53*, to which we have added alterations in *Dicer1* expression. Cre expression, and thus tumorigenesis, is controlled by infecting mice with viral vectors that express Cre, in either a cell type independent (left panel) or cell type specific (middle and right panels) manner. In all models except KP, expression of Cre also deletes one allele of *Dicer1*in those cells. In KPDT-1 mice, we express Cre in Club cells, using an adenoviral vector, and mutate *Dicer1* in ATII cells, using a lentiviral vector. In KPDT-2 mice, we reverse these cell types, expressing Cre in ATII cells and mutating *Dicer1* in Club cells. Mutation of *Dicer1* in a non-tumor bearing cell population accelerates tumor progression and shortens expected survival dramatically in KPDT-1 mice and modestly in KPDT-2 mice.

The highest percentages of animals with pulmonary tumors and lymph node metastases were observed in our KPDS model. In this model, Cre is expressed in club cells under the control of the tamoxifen-inducible *Scgb1a1* Cre driver, and Dicer1 is truncated in AT2 cells. Due to the leakiness of this Cre driver, a small amount of Cre is likely expressed during embryonic development. Comparison of results from KPDS and KPDT-1 animals suggests that tumorigenesis and metastasis are influenced by aging. Additionally, the order of inducing the genetic alterations (expressing *Kras^G12D^,* deleting *Trp53, deleting* one *Dicer1* allele in Club cells, then mutating *Dicer1* in AT2 cells) or the time interval between them may also affect tumorigenesis and metastasis.

We observed a dramatic acceleration of tumor progression only in our KPDT-1 model, in which we expressed Cre in club cells and mutated *Dicer1* in AT2 cells. Mutation of *Dicer1* in AT2 cells in KPDT-1 mice resulted in aggressive adenocarcinomas that surrounded and invaded the airways by ten weeks after viral infection. At twelve weeks after viral infection, KPDT-1 animals began to display signs of respiratory distress, necessitating euthanasia, and their lungs often contained large regions of hemorrhage upon histological examination. KP mice administered the same Ad5CC10Cre viral vector were previously reported to have a median survival of 25.3 weeks (Sutherland et al., 2014), indicating that altering *Dicer1* reduced the expected survival time of these animals by half. We also observed tumor cells invading blood vessels at this time, suggesting that tumor cells in KPDT-1 mice were disseminating through the circulatory system. In the absence of *Dicer1* alteration, pulmonary tumors in KP mice were previously shown to be more invasive and metastatic when initiated in club cells compared to AT2 cells (Sutherland et al., 2014).

Reversing the cell types in KPDT-2 mice by expressing Cre to initiate tumors in AT2 cells and mutating *Dicer1* in club cells resulted in only a modest acceleration of tumor progression. Similarly, we did not observe any effect of Dicer1 mutation in club cells on expected survival, as KPDT-2 animals were harvested at 20 weeks after viral vector administration, and KP mice administered the same Ad5SPCCre viral vector had a previously reported median survival of 23.1 weeks (Sutherland et al., 2014). Using fluorescent microscopy, we observed green fluorescent tumor cells outside of the lungs at 20 weeks after tumor induction, suggesting that tumor cells in KPDT-2 animals can become metastatic if given sufficient time (Fig. S3).

Collectively, these results suggest that LUAD progression is influenced by miRNA expression in a non-cell-autonomous manner, and this potential miRNA-mediated intercellular communication may be unidirectional. In our mouse models, miRNA-mediated intercellular communication may not be direct between club and AT2 cells. Rather, miRNAs could be transferred to an effector cell, such as an immune cell. In a mouse model of acute lung injury, for example, miR-223 is transferred intercellularly from neutrophils to pulmonary epithelial cells (Neudecker et al., 2017). Analysis of transcription in lung tissues from both KPDT-1 and KPDT-2 mice to identify the potential molecular basis of the phenotypic differences reported here will be the focus of future investigations.

Identifying the cancer cell of origin may lead to a better understanding of the earliest events in tumor initiation and the development of therapeutic treatments to interrupt cancer progression (reviewed in (Ferone et al., 2020, Visvader, 2011). Animal studies have shown that LUAD, driven by Kras^G12D^ expression and Trp53 deletion, can arise from multiple cell types (reviewed in (Ferone et al., 2020, Sainz de Aja et al., 2021, Rowbotham and Kim, 2014). In recent studies, AT2 cells have emerged as the presumed cell of origin for LUAD, based primarily upon the aggressiveness, location and histological classification of the resulting tumors. Our results, however, showed that the progression of adenomas to aggressive adenocarcinomas was dramatically accelerated only when tumors were initiated in club cells and *Dicer1* was mutated in AT2 cells. Thus, we assert that club cells can serve as the cell of origin for LUAD in the context of *Dicer1* mutations in AT2 cells. Similarly, a murine model of PPB demonstrated that ablation of *Dicer1* in SPC-positive lung epithelium during development is sufficient to initiate cystic lungs, but not to drive progression of mesenchymal cells to sarcoma (Wagh et al., 2015).

DICER1 has been identified as a substrate of active extracellular signal-regulated kinase (ERK), and phosphorylated nuclear DICER1 functions with oncogenic KRAS to mediate lung tumor progression, independent of epithelial-to-mesenchymal transition (EMT) and miRNA production (Reyes-Castro et al., 2023). We hypothesize that the accelerated tumor progression and reduced expected survival we observed in our KPDT-1 mice are independent of DICER1 phosphorylation. Murine DICER1 protein is phosphorylated at serine 1712 and 1836, and our two *Dicer1* truncating viral vectors cut around amino acid 1687. Analysis of Cas9-targeted sequencing reads of non-tumor epithelial cells predicted that DICER1 would be out of frame in all reads containing indels. Thus, we conclude that the acceleration in tumor progression that we observed in KPDT-1 mice likely occurs through an independent mechanism.

We used the *DICER1* variants present in *DICER1* tumor predisposition as a blueprint for genetic alterations that may affect tumorigenesis. Here, we demonstrate that, in a mouse model, those alterations affect the progression of LUAD, which is not recognized as a DICER1 tumor predisposition tumor type. Thus, DICER1 pathogenic variants could play a wider role in cancer by contributing to the progression of additional tumor types than currently recognized. The prevalence of pathogenic *DICER1* variants in non-cancer databases is estimated to be 1:10,600 (Kim et al., 2017), but escalates to 1:52 in non-small cell lung cancer (NSCLC) cases (Chong et al., 2024). However, to the best of our knowledge, *DICER1* variants have not been examined in non-tumor cells from patients with cancer of the lung or any other organ. While our results clearly demonstrate that *Dicer1* mutations in AT2 cells affect tumors initiated in club cells, we have not tested whether *Dicer1* mutations in other cell types can also influence the progression of such tumors. Since our viral vectors can be modified to express a variety of cell-type-specific promoters, we anticipate that they will be useful in answering many questions about the role of *Dicer1* and miRNA-mediated intercellular communication in tumorigenesis.

## MATERIALS AND METHODS

### Mice

All research involving animals was conducted following Animal Use protocol #01011 approved by the Institutional Animal Care and Use Committee (IACUC) from JAX. Lung cancer models were generated by breeding five different mouse strains, all of which were obtained from The Jackson Laboratory (Bar Harbor, ME): B6.129-*Kras*^tm4Tyj^*Trp53*^tm1Brn^/J (strain #032435, referred to as KP); B6.Cg-*Dicer1*^tm1Bdh^/J (strain #006366, referred to as D); B6N.129S6(Cg)-*Scgb1a1*^tm1(cre/ERT)Blh^/J (strain #016225, referred to as S); B6.129S-*Sftpctm1(cre/ERT2)Blh/J* (strain #028054, referred to as F); and B6.129(Cg)-Gt(ROSA)26Sor^tm4(ACTB-tdTomato,-EGFP)Luo^/J (strain #007676, referred to as T). First, we generated the KPD strain by breeding KP mice to D mice. KPD mice were then bred to S mice to generate KPDS mice and to F mice to generate KPDF mice. An additional strain, (DPT), was generated by breeding D, P, and T mice together. Finally, KPDT mice were generated by breeding the KPT to DPT strains. All models were generated on a C57BL/6J background. The final genotypes of every mouse strain are provided in Table S1.

Mice were genotyped by PCR followed by gel electrophoresis, using the primers listed in Table S2. DNA was isolated from tissues obtained by either ear notching or tail tipping using the Hot Shot method (Truett et al., 2000). Each 20 microliter (μl) PCR reaction consisted of 1 μl of DNA, 1 μl of each genotyping primer (10 millimolar (mM) stock), 4 μl of 5 molar (M) betaine (Sigma-Aldrich, St. Louis, MO), 4 μl of 5X Phusion buffer (New England Biolabs, Beverly, MA), 2 μl of dNTP mix (10 mM each dNTP; Promega, Madison, WI) and 0.2 μl of Phusion DNA polymerase (New England Biolabs, Beverly, MA). Reactions were amplified for the number of PCR cycles at the annealing temperatures indicated in Table S2 and visualized by electrophoresis on a 1.4% agarose gel. The expected sizes of the resulting PCR products are also indicated in Table S2.

### Lentiviral constructs

To create lentiviral constructs expressing Cas9 and guide RNAs, we first identified potential protospacer sequences using Benchling software (benchling.com) and its built-in gene-editing tools. Candidate guides were chosen to overlap with known ‘hotspot’ mutations in *DICER1* and were centered around exons 24 and 25. The protospacer sequence CTCCCAGGAATTCTAAGCGC, referred to as CRISPR guide RNA x24sgRNA1 (mm39; negative strand chr12:104660976-104660995; Eurofins Genomics, Louisville, KY), was chosen for all constructs. The x24Lentiviral plasmids, pLex307iCre, SPCx24 and CC10×24, were constructed by standard molecular biology techniques using parent plasmids obtained from Addgene (Watertown, MA). pLCx24Cas9 was constructed by inserting oligos encoding the CRISPR guide RNA x24sgRNA1 into the BsmBI site downstream of the human U6 promoter in the plasmid pLentiCrispr V2 (Addgene #52961; Watertown, MA). In the resulting plasmid, an ORF encoding iCre (https://pubmed.ncbi.nlm.nih.gov/11835670/) was inserted in place of the puromycin resistance cassette, downstream of the encoded T2A sequence. To create SPCx24 and CC10×24 viral vectors, iCre was deleted from the pLCx24Cas9 plasmid and the plasmid’s EFS promoter was replaced with either the human SPC (hSPC) promoter or the murine CC10 (mCC10) promoter. Promoter fragments were generated by PCR utilizing Q5 High-Fidelity DNA polymerase (New England Biolabs, Beverly, MA) and primers listed in Table S2, following manufacturer’s protocol. For PCR reactions, DNA from mouse strain 006225 (JAX, Bar Harbor, ME) was used to generate an hSPC promoter fragment and from strain 000664 (JAX, Bar Harbor, ME) to generate a mCC10 promoter fragment. pLEX307-iCre was created by inserting the iCre ORF downstream of the EF1 promoter in plasmid pLEX_307 (Addgene #41392, Watertown, MA)

Lentiviral packaging was performed in human 293T/17 cells (ATCC CRL-11268, Manassas, VA) cells using packaging plasmids obtained from Addgene (pMD2.g, Addgene #12259; pMDLg/pRRE, Addgene #12251; pRSV-Rev, Addgene #12253). Lentiviral supernatants were purified using ultracentrifugation through a 20% sucrose cushion, and pellets were resuspended in 1X Hank’s balanced salt solution (HBSS; Mediatech Corning Cellgro, Manassas, VA) prior to aliquoting and storage at -80C. Titration for Cre-containing lentiviral particles was performed on an SV-40 immortalized mT/mG ear fibroblast cell line derived from strain 007676 (JAX, Bar Harbor, ME) and was calculated based on the number of GFP+ cells obtained after infection. Lentiviral particles without Cre or other reporters were titrated in 293T/17 cells using ddPCR with primers/probes against the rev response element (RRE) and the genomic locus RPPH1 (Table S2).

### Single lung cell suspensions

To confirm that our CRISPR guide RNA x24sgRNA induced mutations in *Dicer1* exon 24, we infected a male KPDT mouse with 2.5 x 10^8^ plaque forming units (pfu) of Ad5mSPC-Cre (University of Iowa) and 2.3 x 10^4^ titration units (TU) of CC10×24 by intranasal instillation at 7 weeks of age, made a single cell suspension of his lungs at 26 weeks after infection (Sekiguchi and Hauser, 2019), sorted those cells by FACS and sequenced various cell populations by Cas-9 targeted sequencing (Lesbirel et al., 2021). To make a single lung cell suspension, this mouse was euthanized by CO_2_ asphyxiation and his lungs were perfused with 10 ml of cold 1X Dulbecco’s phosphate buffered saline (DPBS; Gibco Thermo Fisher Scientific, Carlsbad, CA) followed by 3 mls of prewarmed dispase II (1.6 units/ml; Life Technologies, Grand Island, NY)/collagenase type I (300 units/ml; Worthington Lakewood, NJ) in Dulbecco’s Modified Eagle Medium/F12 (DMEM/F12; Life Technologies, Grand Island, NY)(Sekiguchi and Hauser, 2019). All five lung lobes were transferred to a petri dish, DPBS was gently blotted with Kimwipes (Kimberly-Clark, Roswell, GA) and tissues were chopped with a razor blade. Next, 2 mls of dispase (1.6 units/ml)/collagenase (300 units/ml)/DNase I (10 units/ml; Promega, Madison, WI) in DMEM/F12 were added, and tissues were incubated in a humidified 5% CO2 incubator at 37^0^C for 40 minutes, with pipetting up and down after 15 minutes. The dispase was inactivated by adding 2 mls of cold DMEM/F12 plus 5% bovine serum albumin (BSA, Sigma-Aldrich, St. Louis, MO) and tissues were transferred into two 2 ml conical tubes and centrifuged at 500 x g for 5 minutes at room temp. Supernatants were carefully removed, tissue pellets were washed with 2 mls of HBSS (Mediatech Corning Cellgro, Manassas, VA), and combined into one tube. The wash and centrifugation were repeated and 1 ml of protease mix [Accutase (4.5 mls/ml; Innovative Cell Technologies, San Diego, CA), Accumax (4.5 mls/ml; Innovative Cell Technologies, San Diego, CA), Bacillus licheniformis protease (0.1 mg/ml; Creative Enzymes, Shirley, NY) in calcium and magnesium-free DPBS] was added to the resulting tissue pellet. The sample was transferred to a 15 ml conical tube and pipetted up and down with a 1000 ml low-bind pipet tip for 2 minutes. After incubating on ice for 15 minutes, 10 mls of calcium and magnesium-free DPBS plus 10% filtered fetal bovine serum (FBS; R&D Systems, Minneapolis, MA) were added and mixed by pipetting. The sample was passed through a 70 micron nylon cell strainer (Fisher Scientific, Pittsburgh, PA) set over a 50 ml conical tube. The flow-through was centrifuged at 500 x g for 5 minutes at room temp. and the supernatant was carefully removed. The resulting cell pellet was resuspended in 3 mls of red blood cell (RBC) lysis buffer (150 mM ammonium chloride; Sigma-Aldrich, St. Louis, MO, 10 mM sodium bicarbonate; Fisher Scientific, Pittsburgh, PA and 1 mM ethylenediaminetetraacetic acid [EDTA] disodium salt; Sigma-Aldrich, St. Louis, MO) and incubated on ice for 5 minutes. After adding 10 mls of DPBS plus 1% FBS, the sample was centrifuged at 500 x g for 5 minutes at room temp and the supernatant was discarded. The resulting cell pellet was resuspended in 990 mls of calcium and magnesium-free DPBS supplemented with 1% FBS. After adding 1ml pf anti-mouse CD326 (Epcam) antibody (Biolegend, San Diego, CA), the cell sample was incubated on ice, protected from light for 15-20 minutes, then washed twice with 2 mls of DPBS plus 1% filtered FBS. After removing the supernatant, cells were resuspended in 750 mls of DPBS plus 1% filtered FBS and passed through a Flowmi 70 cell strainer (Bel-Art Fisher Scientific, Pittsburgh, PA). Single lung cell suspensions were sorted into three populations, GFP+, tdTomato+ Epcam high and tdTomato+ Epcam low, by using an Attune NXT acoustic focusing cytometer (ThermoFisher Scientific, Waltham, MA).

### Cas9 Targeted Sequencing

Genomic DNA (gDNA) was extracted from GFP+, tdTomato+ Epcam high and tdTomato+ Epcam low cell pellets using the Monarch HMW DNA Extraction Kit (New England Biolabs, Beverly, MA) following the manufacturer’s protocol. CRISPR/Cas9 targeted libraries were constructed as previously described (Gilpatrick et al., 2020) using the two CRISPR RNAs (crRNA; IDT Coralville, IA) listed in Table S2. The resulting capture libraries were sequenced for 24 hours on a GridION using an R9.4.1 flow cell (Oxford Nanopore Technologies, Oxford, U.K.). Base calling was carried out using GUPPY (v3.2.10). The resulting fastq files were aligned to a reference sequence using minimap2(v2.17). Custom reference sequences were constructed using the *Mus* musculus reference genome (GRCm38), and alignment results were subjected to MapQ score filtering using Samtools (v1.11). Subsequent coverage depth for on-target reads was generated using Bedtools (v2.29.2) and was visualized using the Integrative Genomics Viewer (IGV) (Thorvaldsdottir et al., 2013). Consensus sequences were generated using Medaka (v1.0.3).

### Tamoxifen administration

A stock solution of tamoxifen (50 ml/ml; Sigma-Aldrich, St. Louis, MO) was prepared in corn oil (Sigma-Aldrich) and stored at 4^0^C for up to one week. At seven weeks of age, mice were administered 0.04 mls of the stock solution per 10 g of body weight by oral gavage. Mice received either 2 or 4 tamoxifen doses, one dose per day, on consecutive days.

### Viral infection

Mice were administered viral vectors by intranasal instillation, following the protocol of DuPage et al. (DuPage et al., 2009). In brief, five- to seven-week-old mice were anesthetized by intraperitoneal injection of an anesthetic, and Cre-expressing adenovirus or lentivirus was introduced intranasally, followed by *Dicer1*-truncating lentiviral vectors, if used. Mice were allowed to recover on a heated pad, then returned to their cages once they were capable of moving, eating, and drinking. Originally, mice were anesthetized with 2-2-2-Tribromoethanol. However, due to a change in institutional policy, we switched to anesthetizing mice with a mixture of ketamine (4mgs/ml), xylazine (0.4 mgs/ml) and acepromazine (0.1 mg/mls) at a dose of 0.1 mls per 10g of body weight. After administration of virus and viral vectors, mice received 1 ml of warm, sterile saline (VetOne, Kings Park NSW, Australia) per 25 mg of body weight by subcutaneous injection between the shoulder blades, and 0.1 ml per 10 g of body weight of atipamezole (0.1 mg/ml) by intraperitoneal injection. Our selected doses of adenoviruses were 1.0 x 10^9^ pfu/mouse for Ad5mSPC-Cre, and 1.0 x 10^8^ pfu/mouse for Ad5CC10-Cre and Ad5Empty. Adenoviruses were diluted in Dulbecco’s modified Eagle’s medium (DMEM; Gibco, Grand Island, NY) and precipitated by the addition of calcium chloride (Fisher Scientific, Pittsburgh, PA), while lentiviral vectors were diluted in DMEM only. If mice were only receiving adenovirus, the final volume was 50 microliters per animal. If mice received both an adenovirus and a lentiviral vector, the volume per animal was 40 microliters of adenovirus and 10 microliters of lentiviral vector.

### Tissue Collection and Fixation

At the desired time after tumor induction or treatment, mice were euthanized by CO_2_ asphyxiation and lungs were perfused with 10 mls of 1X PBS followed by 5 mls of 4% paraformaldehyde (PFA; Sigma-Aldrich, St. Louis, MO). Organs were carefully removed and fixed in 4% PFA at 4^0^C. Tissue fixation times depended upon the thickness of the organ or tissue and ranged from 1-3 days. After fixation, tissues were rinsed with 1X PBS then embedded in paraffin. Thin sections of each tissue were cut, transferred to glass slides, and stained with hematoxylin and eosin (H&E). All H&E slides of lung tissues were examined and evaluated by a veterinary pathologist (R.D.). To make figures, slides were scanned using a Hamamatsu NanoZoomer slide scanner at 40×.

## Supporting information

Supplemental data, will be used to link to the manuscript file on the preprint site

## ACKNOWLEDGEMENTS

We gratefully acknowledge the contributions of the Histopathology & Microscopy, Flow Cytometry and Viral Vector Core Services at The Jackson Laboratory for expert assistance with the work described in this publication. We are also indebted to the dedicated animal care team at The Jackson Laboratory. Finally, we would like to thank Dr. Carla F. Kim for critical reading and helpful discussions.

## AUTHOR CONTRIBUTIONS

Conceptualization: J.W., R.S.M.; Methodology: J.W., R.S.M.; Investigation: J.W., R.D., A.T., W.M., T.M., N.M.H., P.J.K.R., S.L., J.R.C., H.M., T.B.; Resources: R.S.M., Writing-original draft: J.W.; Writing-review & editing: R.S.M., C.J.B.; Supervision: J.W.; Project Administration: C.J.B.; Funding acquisition: J.W., C.J.B.

