## Supplemental data, will be used to link to the manuscript file on the preprint site for "Cell-type Specific Alteration of *Dicer1* Accelerates Tumor Progression in Mouse Models of KRAS-driven Lung Adenocarcinoma"

### SUPPLEMENTAL MATERIALS AND METHODS

#### Optimizing Ad5CC10 and Ad5mSPC dosing

We optimized the amounts of Ad5CC10-Cre and Ad5mSPC-Cre viruses for infection of the targeted cell populations by counting GFP+ cells following intranasal instillation of one of three doses of either virus (Fig. S2A). Cohorts (n=5) of six- to eight-week-old KPDT mice were anesthetized and intranasally administered 1.0 x $10^8$ pfu, 5.0 x $10^8$ pfu or 2.5 x $10^9$ pfu of Ad5SPC-Cre. Additional cohorts of mice were intranasally administered 1.0 x $10^7$ pfu, 5.0 x $10^7$ pfu or 2.5 x $10^8$ pfu of Ad5CC10-Cre. Fifteen days post viral infection, all animals were euthanized and their lungs were harvested. Single cell suspensions of lung tissues from four animals of each cohort were analyzed on a FACSymphony A5 (BD Biosciences Franklin Lakes, NJ) to count the numbers of TdTomato$^+$ and GFP$^+$ cells.

Upon examining the frequency of GFP+ cells in the lungs of all six cohorts, including cells positive for both tdTomato and GFP, we found a statistically significant difference (Mann-Whitney test; p=0.0286) between the low and mid doses for both viruses but not between the mid and the high doses (Mann-Whitney test; p=0.2000 for Ad5CC10-Cre and p=0.1143 for Ad5mSPC-Cre, Fig.S2A). The median frequencies of GFP+ cells for mice treated with Ad5CC10-Cre ranged from 0.007165% (low dose) to 0.1155% (high dose). The median frequencies of GFP+ cells for mice treated with Ad5mSPC-Cre ranged from 0.0001595% (low dose) to 0.01935% (high dose).

Lungs tissues from the fifth animal of each cohort were fixed for fluorescent microscopy as described below. Slides were stained with DAPI (1 μg/ml; Thermo Scientific, Waltham, MA) and scanned on a Leica Thunder 4-slide scanner (Leica Microsystems, Wetslar, Germany). Fluorescent microscopy examination of these tissues confirmed that these viruses were infecting the intended cell populations (Fig. S2B).

#### Fluorescent Microscopy

Tissues for fluorescent microscopy were harvested in the absence of bright light. After fixation in 4% PFA, organs were transferred to a series of sucrose (Fisher, Pittsburgh, PA) solutions, 10%, 15%, 20% and 30%, and remained in each solution until tissues sank to the bottom of the 50 ml conical tubes. When ready, tissues were removed from the final sucrose solution, gently blotted on a Kimwipe then transferred to a cryomold, embedded in Tissue Tek Optimal Cutting Temperature (OCT; Sakura, Torrance, CA) and stored at -80$^0$C. Tissues were cut into 30-micron thick sections using a cryostat, transferred to slides and stored at -80$^0$C. To stain, slides were incubated in the dark at room temperature for 60 minutes, then washed three times in 1X PBS. Tissues were covered with DAPI (1 μg/ml; Thermo Scientific, Waltham, MA) and incubated at room temperature for 2-10 minutes, protected from light. Slides were then washed three times with 1X PBS, followed by once with sterile water. The side of each slide was gently touched to a paper towel to remove excess liquid then tissues were covered with Vectashield antifade mounting medium (Vector Laboratories, Burlingame, CA). A coverslip was placed over each tissue and sealed with nail polish. Slides were visualized on a Nikon Ti Eclipse microscope and photographed in the appropriate color channel. The presence of the mT/mG dual fluorescence Cre reporter allowed us to visualize metastatic GFP$^{+}$ cells in tissues outside of the lungs in our animal models (Fig. S3).

**Table S1 Genotypes of mouse strains used in this study**

| Strain | floxed *Kras*$^{G12D}$ | floxed *Trp53* | floxed *Dicer1* | mT/mG | *Scgb1a1*$^{cre/ERT}$ | *Sftpc*$^{cre/ERT2}$ |
|---|---|---|---|---|---|---|
| KP | het | hom | N/A | N/A | N/A | N/A |
| KPD | het | hom | het | N/A | N/A | N/A |
| KPT | het | hom | N/A | hom | N/A | N/A |
| KPDS | het | hom | het | N/A | het | N/A |
| KPDF | het | hom | het | N/A | N/A | het |
| KPDT | het | hom | het | het | N/A | N/A |
| DPT | N/A | hom | hom | hom | N/A | N/A |

**Table S2 Sequences of primers and probes**

| Gene | Forward Primer | Reverse Primer | $T_m$ | PCR cycles | WT band | Mut band |
|---|---|---|---|---|---|---|
| Kras | Kras WT F<br>5'-TGTCTTTCCCCAGCACAGT-3' | Kras Common<br>5'-CTGCATAGTACGCTATACCCTGT-3' | 64$^0$C | 28 | 250 bp | |
| Kras | Kras Mut F<br>5'-GCAGGTCGAGGGACCTAATA-3' | Kras Common<br>5'-CTGCATAGTACGCTATACCCTGT-3' | 64$^0$C | 28 | | 100 bp |
| Trp53 | p53 exon 10 Forward<br>5'-AAGGGGTATGAGGGACAAGG-3' | p53 exon 10 Reverse<br>5'-GAAGACAGAAAAGGGGAGGG-3' | 62$^0$C | 30 | 430 bp | 584 bp |
| Dicer1 | Dicer 6305<br>5'-CCTGACAGTGACGGTCCAAAG-3' | Dicer 6569<br>5'-CATGACTCTTCAACTCAAACT-3' | 56$^0$C | 32 | 351 bp | 420 bp |
| mT/mG | Gt(ROSA) Common<br>5'-AAGGGAGCTGCAGTGGAGTA-3' | Gt(ROSA) WT R<br>5'-GATGACTACCTATCCTCCCA-3' | 62$^0$C | 30 | 357 bp | |
| mT/mG | Gt(ROSA) Common | Gt(ROSA) Mut R | 62$^0$C | 30 | | 284 bp |

| | | | | | | |
|---|---|---|---|---|---|---|
| | 5'-AAGGGAGCTGCAGTGGAGTA-3' | 5'-CGGGCCATTTACCGTAAGTTAT-3' | | | | |
| ßSftpc | Sftpc CM<br>5'-TGCTTCACAGGGTCGGTAG-3' | Sftpc WT R<br>5'-CATTACCTGGGGTAGGACCA-3' | 65$^{0}$C | 31 | 327 bp | |
| Sftpc | Sftpc CM<br>5'-TGCTTCACAGGGTCGGTAG-3 | SftpcMut R<br>5'-ACACCGGCCTTATTCCAAG-3' | 65$^{0}$C | 31 | | 210 bp |
| Scgb1a1 | Scgb1a1 Common<br>5'-ACTCACTATTGGGGGTGTGG-3' | Scgb1a1 WT R<br>5'-AGGCTCCTGGCTGGAATAGT-3' | 62$^{0}$C | 30 | 550 bp | |
| Scgb1a1 | Scgb1a1 Mut F<br>5'-TGATAAGTCTCTGCACCACTACTG-3' | Scgb1a1 Mut R<br>5'-TCGTCAAGAAGACAGGGCCAG-3' | 70$^{0}$C | 28 | | 345 bp |
| RRE | RRE Forward<br>5-TGTGCCTTGGAATGCTAGT-3' | RRE Reverse<br>5'-AATTTCTCTGTCCCACTCCATC-3' | 60$^{0}$C | 40 | 85 bp | |
| RPPH1 | RPPH1 Forward<br>5'-CGCGCGAGGTCAGACT-3' | RPPH1 Reverse<br>5'-GGTACCTCACCTCAGCCATT-3' | 60$^{0}$C | 40 | 135 bp | |
| | **Cas9 Targeted Sequencing** | | | | | |
| Dicer1 | 5'crRNA<br>5'-GGACAATTGTGTAAGTGTCG-3' | 3'crRNA<br>5'-AGCTCTGTCCTGATACCTAG-3' | | | | |

| Probe | Sequence |
|---|---|
| RRE | 5'FAM-TTTGGAATCACACGACCT-3'BHQ |
| RPPH1 | 5'HEX-CCGGCGGATGCCTCCTT-3'BHQ |

**Fig. S1 Detection of leaky Cre expression in KPDS and KPDF mice.** (A) PCR analysis of DNA obtained from the tails of an untreated wild-type and an untreated KPDF mouse, the lung tumors and the tails of three KPDF mice that received corn oil only. The wild-type *Kras* allele generates a band of 250 bps, while the LSL allele of *Kras* generates a band of 100 bps before Cre mediated recombination and 290 bps after recombination. (B) PCR analysis of DNA obtained from the tails of a KPDS mouse that received tamoxifen, an untreated wild-type C57BL/6J mouse, and the lung tumors and tails of three KPDS mice that did not receive tamoxifen. The wild-type *Kras* allele generates a band of 294 bps, while the LSL allele of *Kras* generates a band of 144 bps before Cre mediated recombination and 334 bps after recombination.

**Fig. S2 Optimization of viral tumor induction.** (A) Frequency of GFP$^+$ cells determined by flow cytometry in KPDT mice administered low, mid or high doses of Ad5mSPC-Cre (left panel) or Ad5CC10-Cre (right panel). Virus doses for Ad5mSPC-Cre were1.0 x $10^8$ pfu (low), 5.0 x $10^8$ pfu (mid) or 2.5 x $10^9$ pfu (high) and for Ad5CC10-Cre were 1.0 x 107 pfu (low), 2.5 x $10^8$ pfu (mid) and 2.5 x $10^8$ (high). Significance (*) is p<0.05. (B) Fluorescent microscopy images of lung tissues from KPDT mice infected with a high dose of either Ad5mSPC-Cre (left panel) or Ad5CC10-Cre (right panel) show that the intended cell populations express GFP. Top row images show tissues at lower magnification and bottom row images show boxed regions at higher magnification.

**Fig. S3 Visualization of tumors and metastatic cells in KPDT mice.** Fluorescent microscopy images of tissues from KPDT mice collected in either the DAPI (first column), TdTomato (second column), GFP (third column) or composite (fourth column) channels. All tissues were collected at 20 weeks after viral infection. (A) Tumor cells in KPDT-2 mouse lungs express GFP, but not TdTomato, fluorescently marking them. Both tumor and non-tumor cells show nuclear DAPI staining. (B) Cells in lung tissues from KPDT-2Empty mice express TdTomato, but not GFP. (C) Detection of GFP positive metastatic cells (white arrow heads) in the connective tissue encasing the mediastinal lymph nodes above the heart and behind the thymus from a KPDT-2 mouse.

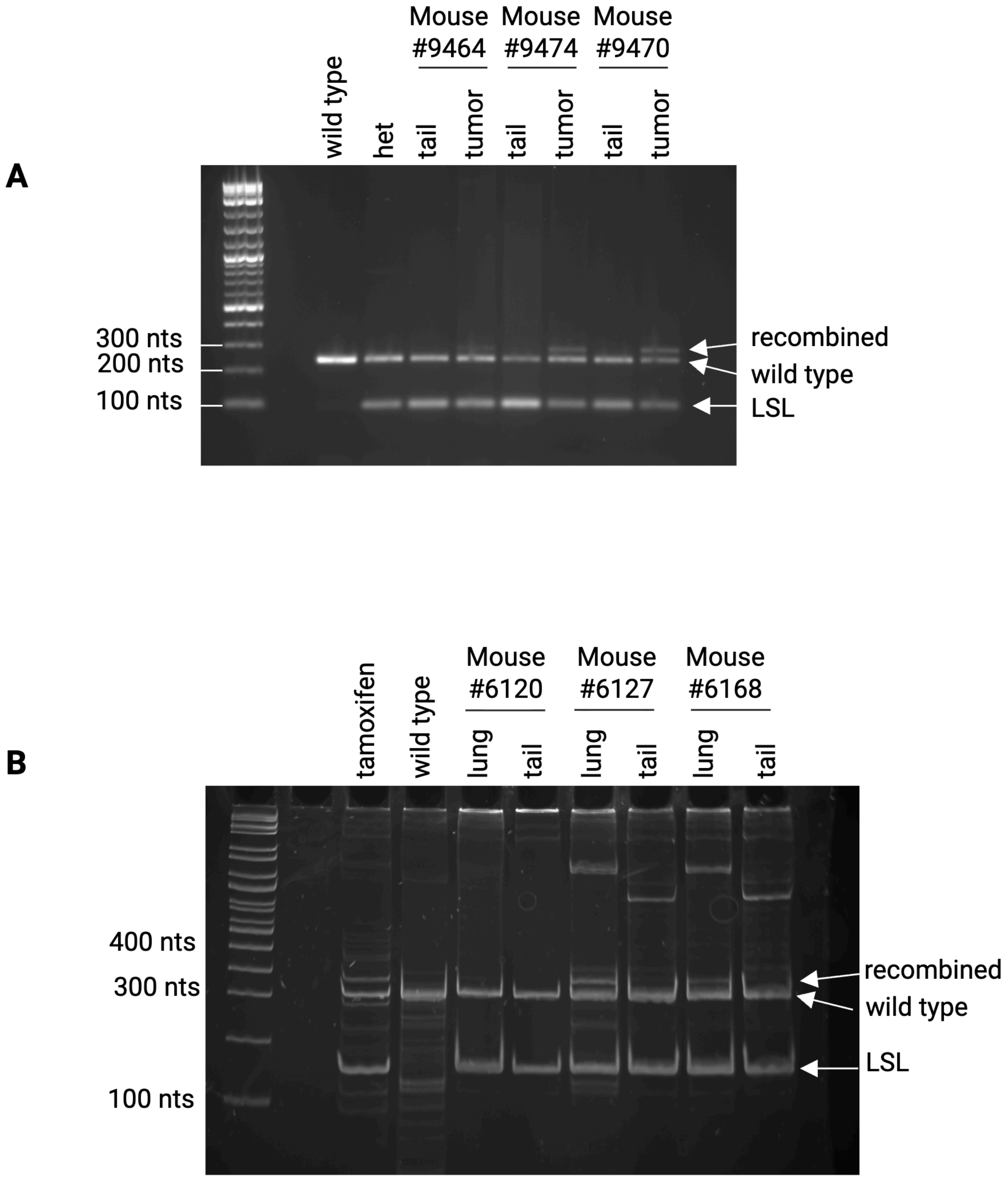
A
Mouse #9464
Mouse #9474
Mouse #9470
wild type
het
tail
tumor
tail
tumor
tail
tumor
300 nts
200 nts
100 nts
recombined
wild type
LSL
B
Mouse #6120
Mouse #6127
Mouse #6168
tamoxifen
wild type
lung
tail
lung
tail
lung
tail
400 nts
300 nts
100 nts
recombined
wild type
LSL

Fig. S1

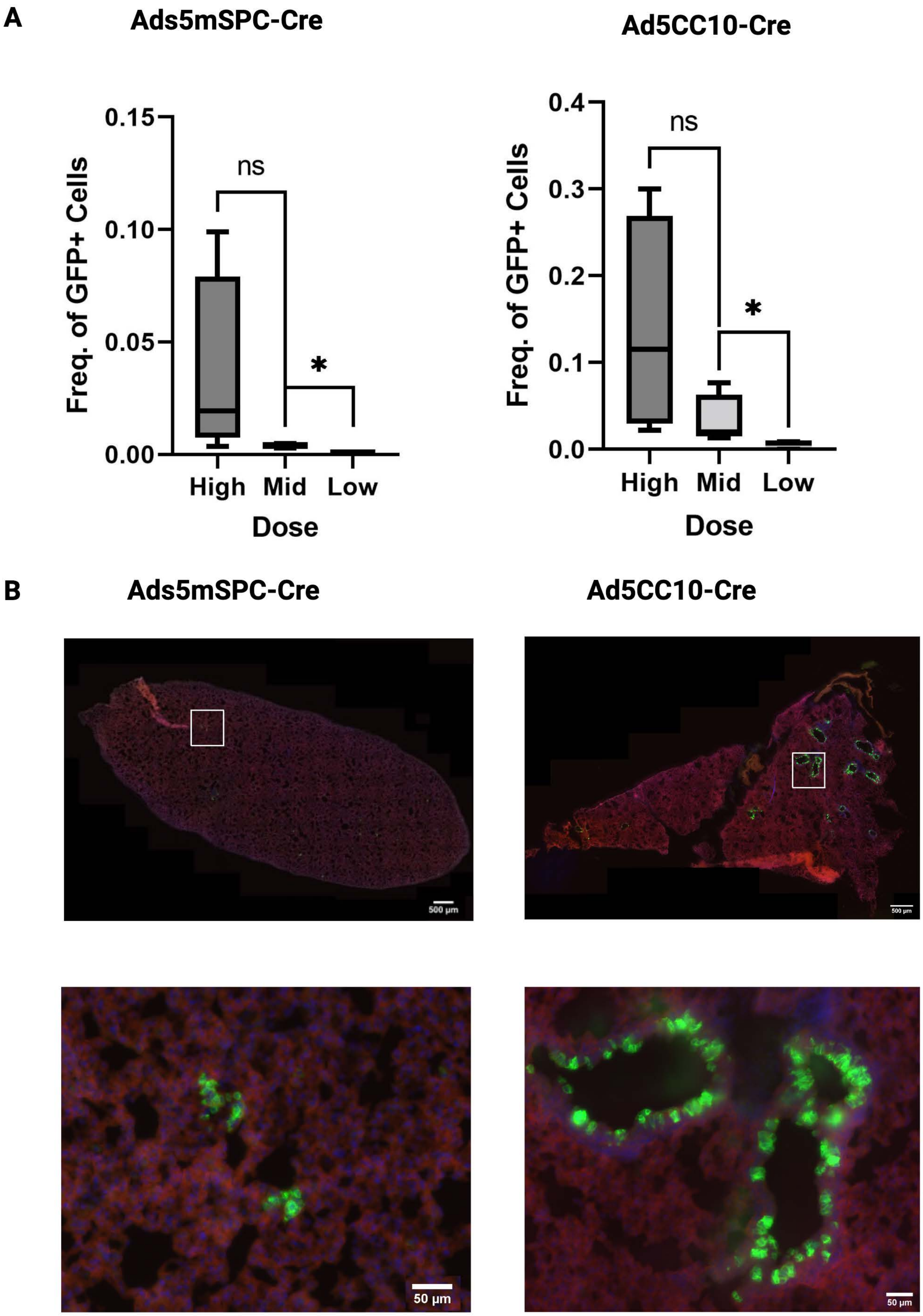
A
Ads5mSPC-Cre
Ad5CC10-Cre
0.15
0.10
0.05
0.00
Freq. of GFP+ Cells
ns
*
High
Mid
Low
Dose
0.4
0.3
0.2
0.1
0.0
B
Ads5mSPC-Cre
Ad5CC10-Cre
500 µm
50 µm

Fig. S2

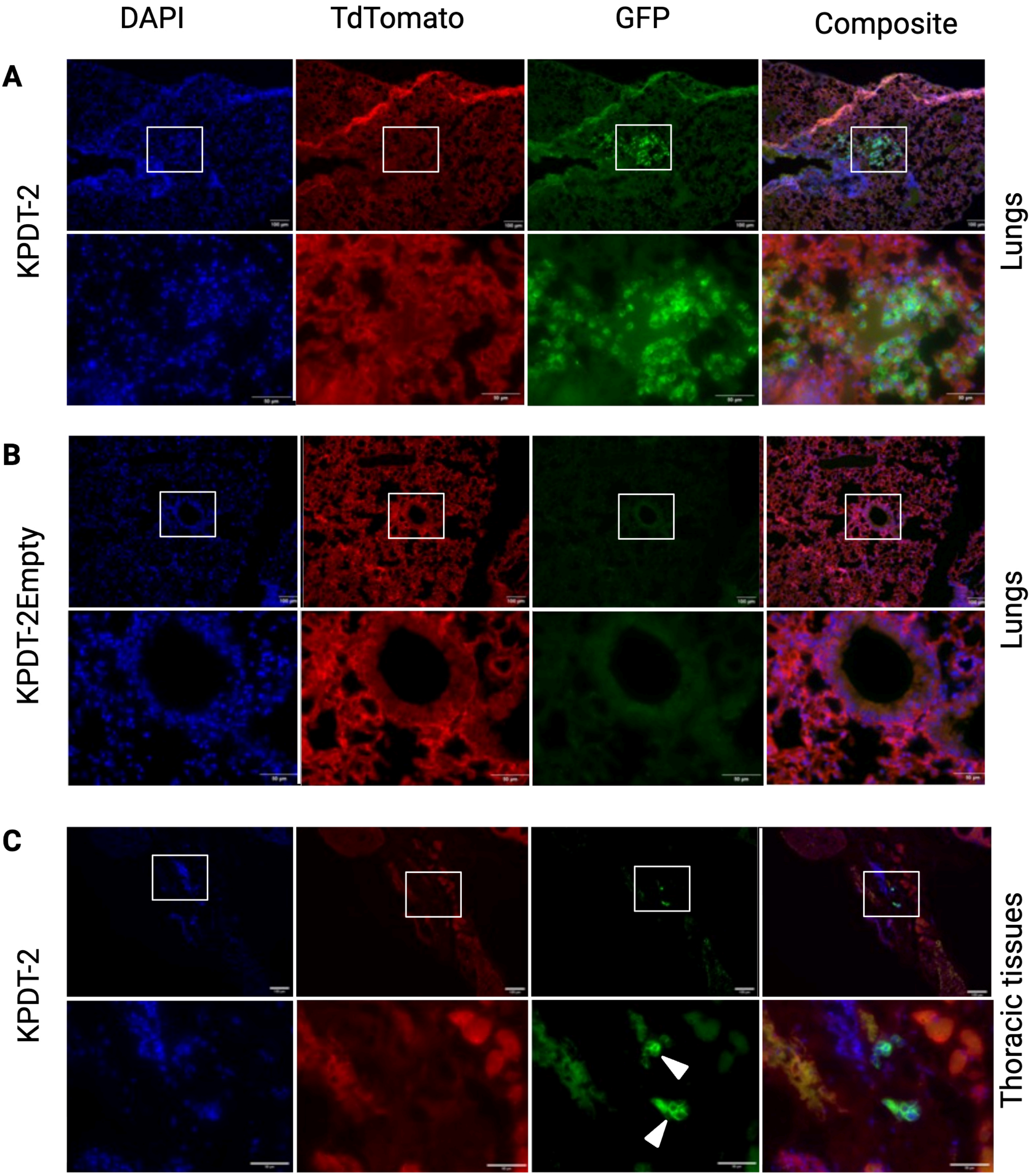
DAPI
TdTomato
GFP
Composite
A
KPDT-2
Lungs
B
KPDT-2Empty
Lungs
C
KPDT-2
Thoracic tissues

Fig. S3
